# Graph in Graph (GiG): A novel graph AI framework for integrating and interpreting medical and omics data

**DOI:** 10.64898/2026.07.20.739657

**Authors:** Heming Zhang, Yifei Lu, Kaiwen Fang, Zixi Xu, Vaha Akabry Moghaddam, Ping An, Shiow Jin, Mary Wojczynski, Michael Province, Fuhai Li

## Abstract

Medical records and omics data are rapidly becoming standard in healthcare settings, which characterize the whole-person from dysfunctional molecules to phenotypes, and thus offer potential for precise disease diagnosis and target discovery. Whereas, it remains an open problem to systematically integrate and interprete medical record and omics data of individual patients. In this study, for the first time, we propose a novel graph AI model framework, Graph in Graph (GiG), to integrate and interpret the whole-person medical and omic datasets. Specifically, the medical record data is modeled using a person-phenotype graph, followed by omics signaling graphs of invidival patients, which enables the integration of information learned from omic-signaling graph and medical phenotype features to characterize individual patients and to prioritize important omic biomarkers and phenotypes. As an exploratory study, we applied and evaluated the GiG model to study the type 2 diabetes (T2D) and pre-T2D vs healthy using the Long Life Family Study (LLFS) cohort, which enrolls families with exceptional longevity to uncover biological mechanisms of healthy aging with medical and omics data. The evaluation results showed that GiG not only achieve a high prediction but also can interpret the prediction by ranking the essential clinical and omic biomarkers. The GiG framework can be applied to other studies by effectively integrating and interpreting medical and omic datasets for disease diagnosis and pathogenesis discovery.

## 1. Introduction

The availability of both medical records and omics data is rapidly becoming standard in healthcare settings, offering immense potential for precise disease diagnosis and target discovery. However, integrating these distinct modalities presents unique computational hurdles. Medical phenotypic data are typically sparse and consist of mixed data types, making it difficult to align patients or accurately estimate phenotypic similarities across diverse clinical conditions. Conversely, omics data are high-dimensional, capturing complex signaling pathways and molecular functions. While integration of these datasets can holistically characterize individual patients to identify key phenotypic factors and genetic biomarkers, systematic integration and interpretation remain in their early stages and an open problem^1,2^. Thus, novel AI models are required to mitigate these new and unique data challenges^1,2^.

The National Institution of Aging (NIA) Long Life Family Study (LLFS)^3^ project was established in 2005 to study EL and systematically identify the genetic and environmental factors that contribute to exceptional longevity. LLFS is an expansive study spanning multiple centers and nations, focusing on the genetic and environmental factors contributing to extraordinary longevity and healthy aging. LLFS recruited families with exceptional longevity, providing a unique resource to identify protective factors for healthy aging and diverse chronic diseases, like type 2 diabetes (T2D), pre-T2D and alzheimer’s diseases. In LLFS, comprehensive data are collected including both medical phenotypes, like demographic information, physical measurements, cognitive abilities, daily activities, and various health markers, and multi-omics data from blood samples. In the LLFS project, one challenge is to integrate and interpret the phenotypic (demographic and clinical) data and omic data to identify the key factors and biomarkers associated with EL.

Artificial intelligence (AI) models have been widely used in the healthcare field^4–16^. Especially, the Graph Neural Networks (GNNs)^17–23^, have revolutionized the field of learning with graph-structured data and empirically achieved the current state-of-art performance in various graph learning tasks, including node classification, link prediction, and graph classification^24^. Broadly, GNNs follow a recursive neighborhood aggregation scheme where the node features from the neighborhood of each node are aggregated to update the node’s feature. Such framework allows GNNs, as one of the most prominent AI models, to be more interpretable compared with other non-GNN AI models^25–28^. The GNNs have been widely used in modeling signaling pathways with omics^14,29–35^, based on three advantages: 1) They can naturally convert graph-structured data into a numerical space, adeptly handling the sparse and mixed data types of phenotypic data; 2) They model omics data in a biologically meaningful way by mapping this data onto signaling pathways or protein-protein interactions; 3) They facilitate the identification of crucial protective factors, which can be framed as the interpretable target-ranking and subnetwork discovery challenges tackled by GNNs through attention analysis. These capabilities are further manifested in their applications in biomedicine, where GNNs are reshaping our comprehension of diseases by elucidating the intricate connections among genes and cells.

In this study, for the first time, we presented a novel deep learning framework, ***Graph in Graph (GiG phenotype-genotype network)***, to resolve the aforementioned data analysis challenges in integrating and interpreting medical phenotype and omics datasets. Specifically, we first convert data frame format phenotypic data into a phenotype graph/network, which links individual T2D patients to phenotypes. Also, each patient will contain an embedded genotype graph in the constructed phenotype network. Thus, all phenotypic nodes and genotypes for each patient can be used as the input of an attention-based model (GAT^19^ and Graph Transformer^18^) to predict the types of individual patients. The advantage of using an attention-based model is that the attention scores of the phenotypes can indicate the importance of phenotypes and gene biomarkers, which makes the model interpretable and facilitates the clinical decision-making of longevity. In the following sections, the data analysis results and details of the methodology are presented.

## 2. Results

### 2.1 GiG architecture and analysis workflow overview

Figure 1a illustrates the architecture of the GiG framework, which employs a dual-graph transformer mechanism to integrate gene-level and patient-level features for classification (Section 4.3). As shown in Figure 1b, we further established a downstream validation pipeline to link the top 142 prioritized genes to pharmacological evidence. By cross-referencing these genes with ChEMBL and conducting citation-consistency checks, we generated a curated ledger of candidate target-drug pairs, which was then overlaid onto the GiG core network (Section 4.5).

**Figure 1.**
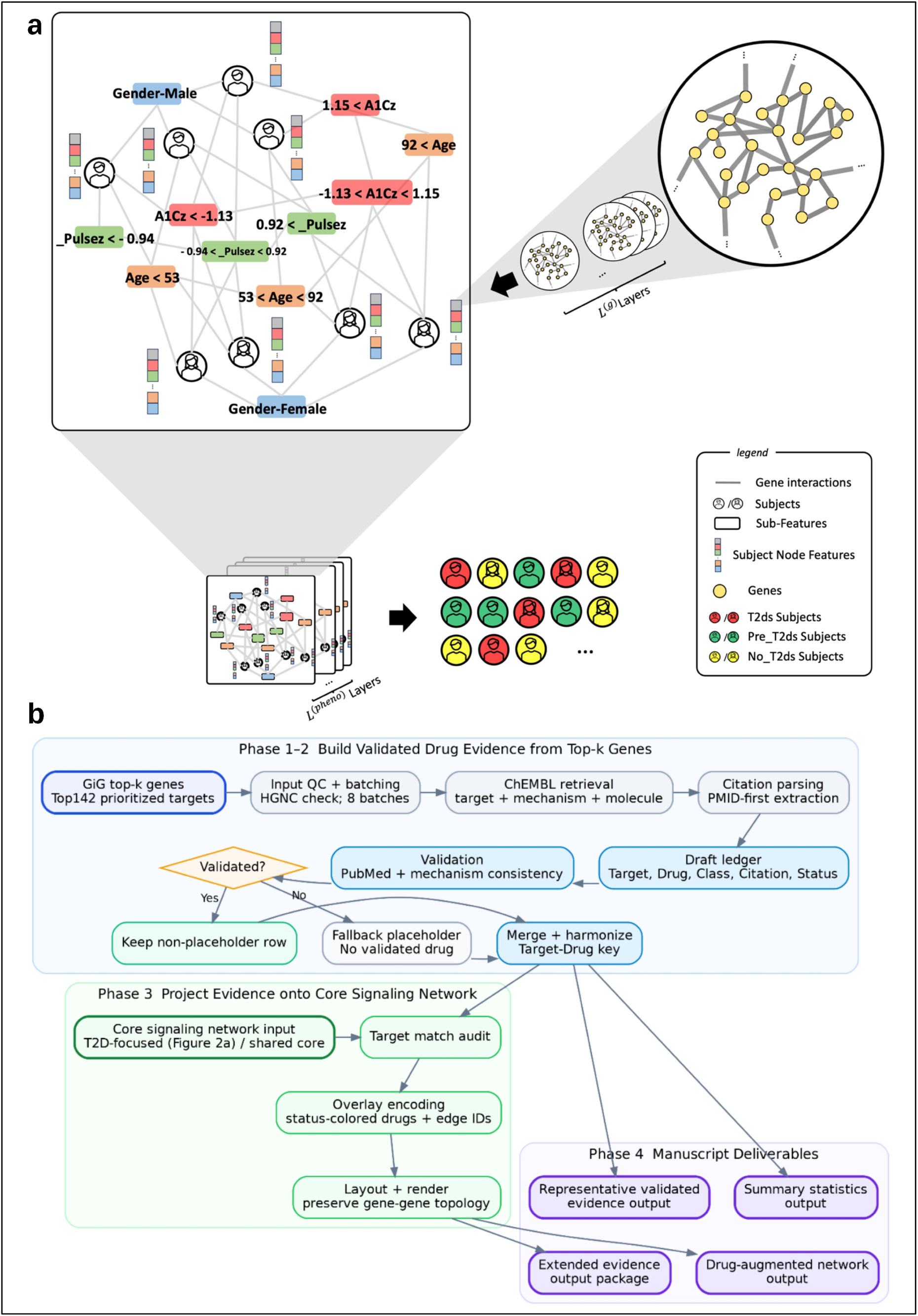
**a.** Overview of the GiG Model Architecture for T2D Risk Stratification. **b.** Workflow for post-GiG drug-evidence augmentation. GiG top-k genes undergo ChEMBL retrieval, PMID-first parsing, and PubMed/mechanism validation, then are harmonized into a target-drug evidence key, projected onto the core signaling network, and reported as summary statistics, representative validated evidence, a drug-augmented network, and an extended evidence package.

### 2.2 Identification of Key Biomarkers for Pre-T2D and T2D via the GiG Framework

This study collected 813 patients’ electronic records with 122 sub-features from 41 characteristic groups (Table 1). Based on the classification, patients have been divided into three categories: 104 pre_t2ds, 68 t2ds, and 641 no_t2ds.

**Table 1.** 41 Characteristic Groups.

|  |  |  |
| --- | --- | --- |
| Sex | Age | HAI |
| FEV1 (LK) | FEV1 | FEV6 |
| FEV1/FVC | FVC | PP-FEV1 |
| PP-FEV6 | BMI | Waist Circ. |
| Body Weight | Arm Span | Pulse Rate |
| SBP | DBP | Pulse Pressure |
| ABI | MMSE | Animal Fluency |
| Digit Forward | Digit Backward | Digit Symbol |
| Logical Mem. (Imm.) | Logical Mem. (Del.) | Total Score (SPPB) |
| Gait Speed | 5x Stand Test | Grip Strength |
| Albumin | Cholesterol | Creatinine |
| Cystatin C | hsCRP | NT-BNP |
| Transferrin | Inorg. T (TSadj) | TSadj BP |
| D2 | D3 |  |

We applied GiG to analyze the medical phenotype and omics data of Pre_T2D and T2D. With the trained parameters of GiG in the transformer, we can obtain the edge weight for the genetic network. We also developed a tool for visualizing the critical genes and components in different groups of patients. By selecting the edge weight threshold, we can filter out the edges and genes to shrink the network to the core graph, indicating the critical signaling pathways and genes (check **Figure 2**).

**Figure 2.**
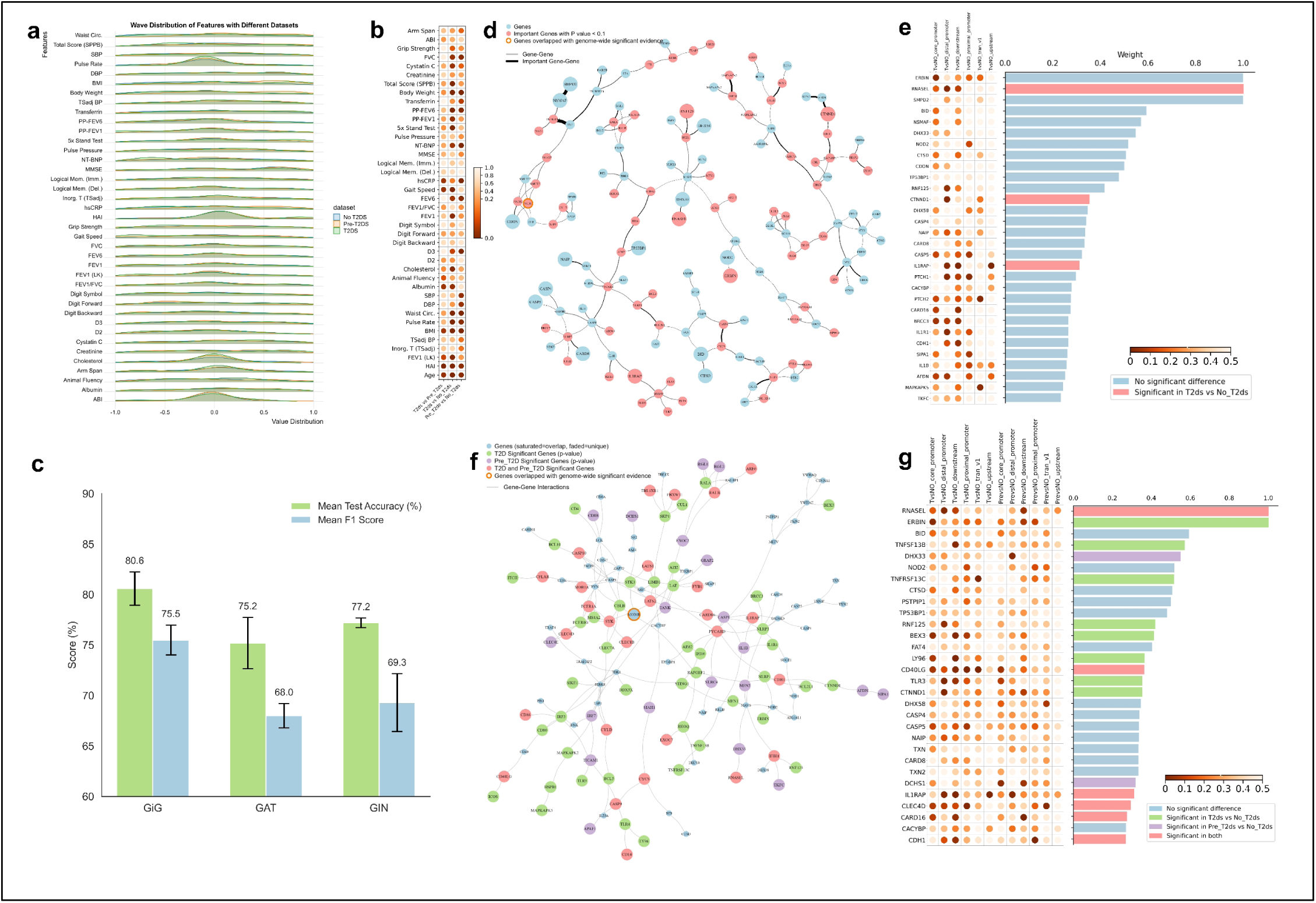
Statistical analysis of clinical phenotypes. **a.** illustrative of wave distribution of features in each clinical phenotypes for each type of patients (T2ds, Pre_t2ds, No_t2ds). **b.** the two-sample Kolmogorov-Smirnov test was demonstrated by comparing the three different types of patients in each clinical phenotype. **c.**Performance comparison between the full GiG model and baseline GNN architectures. Mean test accuracy and mean F1 score are shown for the full GiG model using both phenotype and genotype data and the baseline GAT and GIN models. Error bars represent the standard deviation across cross-validation folds. **d.** the visualization of the top selected nodes by filtering out of the edge weight lower than 0.05 in t2ds patient cohort. The red nodes represent the nodes with any p-value lower than 0.05 when comparing the gene expression and methylation in t2ds cohort and no_t2ds cohort. **e.** the bar plot of the node weights for all of top selected genes (red bar are the genes with any p-value lower than 0.05 by comparing the gene expression and methylation in t2ds cohort and no_t2ds cohort) and the left panel of the bar plot shows the p-value in each type of comparison. **f.** the visualization of the top selected genes combining all of the genes filtered out of the edge weight lower than 0.13 in t2ds patient cohort and the genes filtered out of the edge weight lower than 0.13 in pre_t2ds patient cohort. The green, blue nodes represent the nodes with any p-value lower than 0.05 when comparing the gene expression and methylation in t2ds cohort and no_t2ds cohort and the nodes with any p-value lower than 0.01 when comparing the gene expression and methylation in pre_t2ds cohort and no_t2ds cohort respectively, and the red nodes are the overlapped genes for above 2 types of comparisons. **g.** the bar plot demonstrates the nodes importance in different type of nodes (the colors align with the visualization in the network), and the bottom panel shows the p-value in each type of comparison

Of the top 50 highest-weighted genes identified by the model, 48 have documented associations with T2D or closely related metabolic and inflammatory processes in the published literature, providing broad empirical support for the biological relevance of the model’s output. The most prominent functional cluster, comprising top-weighted genes such as DHX33, CARD8, BRCC3, CASP4/CASP5, and APAF1, centers on pyroptosis and NLRP3 inflammasome signaling, a pathway with well-established mechanistic links to T2D pathogenesis^36^. DHX33 has been shown to sense cytosolic RNA and activate the NLRP3 inflammasome^37^, while CARD8 harbors genetic variants that modulate inflammasome-associated diabetes risk^38^. BRCC3 regulates NLRP3 activity through deubiquitination^39^, and CASP4/CASP5 are activated by saturated fatty acids to trigger downstream IL-1β and IL-18 release^40^. APAF1 has further been implicated in pyroptosis-related retinal pathology in diabetic contexts^41^. Taken together, these genes converge on a coherent inflammatory axis that is mechanistically consistent with β-cell dysfunction and systemic insulin resistance in T2D.

The second functional cluster couples β-cell and insulin signaling with innate immune activation. ERBIN suppresses NOD2-dependent NF-κ B activation and inflammasome-mediated innate immune signaling, and is downregulated in diabetic tissue^42^, and IL1RAP, as the obligate co-receptor of the IL-1 receptor complex, mediates IL-1 β -driven NF-κ B activation and β -cell dysfunction in pancreatic islets^43^, while TXNIP methylation has been independently associated with both inflammation and diabetes^44^. These findings are complemented by innate immune genes with established T2D relevance: NOD2 deficiency aggravates T2D-associated metainflammation^45^, TLR3 displays inflammatory imbalance in diabetic patients^46^, STING1 has been proposed as a therapeutic target in T2D^47^, and IFIH1 variants associate with both autoimmune and type 2 forms of diabetes^48^.

Among the 50 genes, includes high-confidence hits such as TNFRSF13C^49^ and CD40LG^50^. The convergence between model-assigned weights and independently reported genetic and functional evidence indicates that the node scoring scheme captures biologically meaningful disease signal rather than statistical noise.

### 2.3 Pre_T2D and T2D stage-specific targets and pathways

To further characterize the biological relevance of the GiG network, we examined differential RNA-seq expression and disease-stage specificity among all 142 high-weight genes (the full set from which the top 50 were selected) identified by the model. Figure 2a presents the disease-stage specificity landscape of these genes, plotting the significance of regulatory-region methylation differences (Mann–Whitney U test; minimum raw p-value across six regulatory regions per gene) in T2D versus No-T2D (x-axis) against Pre-T2D versus No-T2D (y-axis), with bubble size proportional to network edge weight. The genes partition into three distinct groups: 25 genes reach significance in both comparisons (both significant), reflecting dysregulation shared across early and established disease; 45 genes are T2D-specific, achieving significance only against the T2D versus No-T2D axis; and 19 genes are Pre-T2D-specific, elevated in significance on the Pre-T2D axis alone. Among the most prominent both-significant genes, CARD16, CDH1, and IFIH1 occupy the upper-right quadrant with high network weights, suggesting that these genes are not only strongly co-expressed within the GiG network but also consistently dysregulated from the pre-diabetic stage onward. IL1B and SIPA1 likewise fall in this quadrant with moderate network weights, consistent with their established roles in inflammasome signaling^51^ and immune cell migration^52^. Conversely, T2D-specific genes such as IL1RAP, RNASEL, and LTB, which carry some of the highest x-axis significance scores, may reflect molecular changes that emerge only in frank disease rather than during the metabolic transition preceding it.

Panel A of Figure 3b strengthens these network-level findings at the individual gene level by displaying residual RNA-seq expression distributions for four representative high-weight genes across the three disease groups. IRF7 exhibits a significant and progressive decline from No T2D to T2D (FDR < 0.01, Cohen’s d = −0.45), indicating that interferon regulatory factor 7-mediated innate immune signaling is suppressed as disease advances^53^. CD40LG, CYLD, and TNFRSF13C each show a trend toward higher expression in T2D relative to No T2D (d = +0.37, +0.38, and +0.48, respectively; FDR < 0.10), pointing to modest but consistent upregulation of co-stimulatory^54^, deubiquitinase^55^, and B-cell survival signaling in the T2D state. Taken together, these expression patterns corroborate the network weights assigned by the GiG model, reinforcing the interpretation that the top-ranked genes capture genuine disease-associated transcriptional signals rather than statistical artifacts.

**Figure 3.**
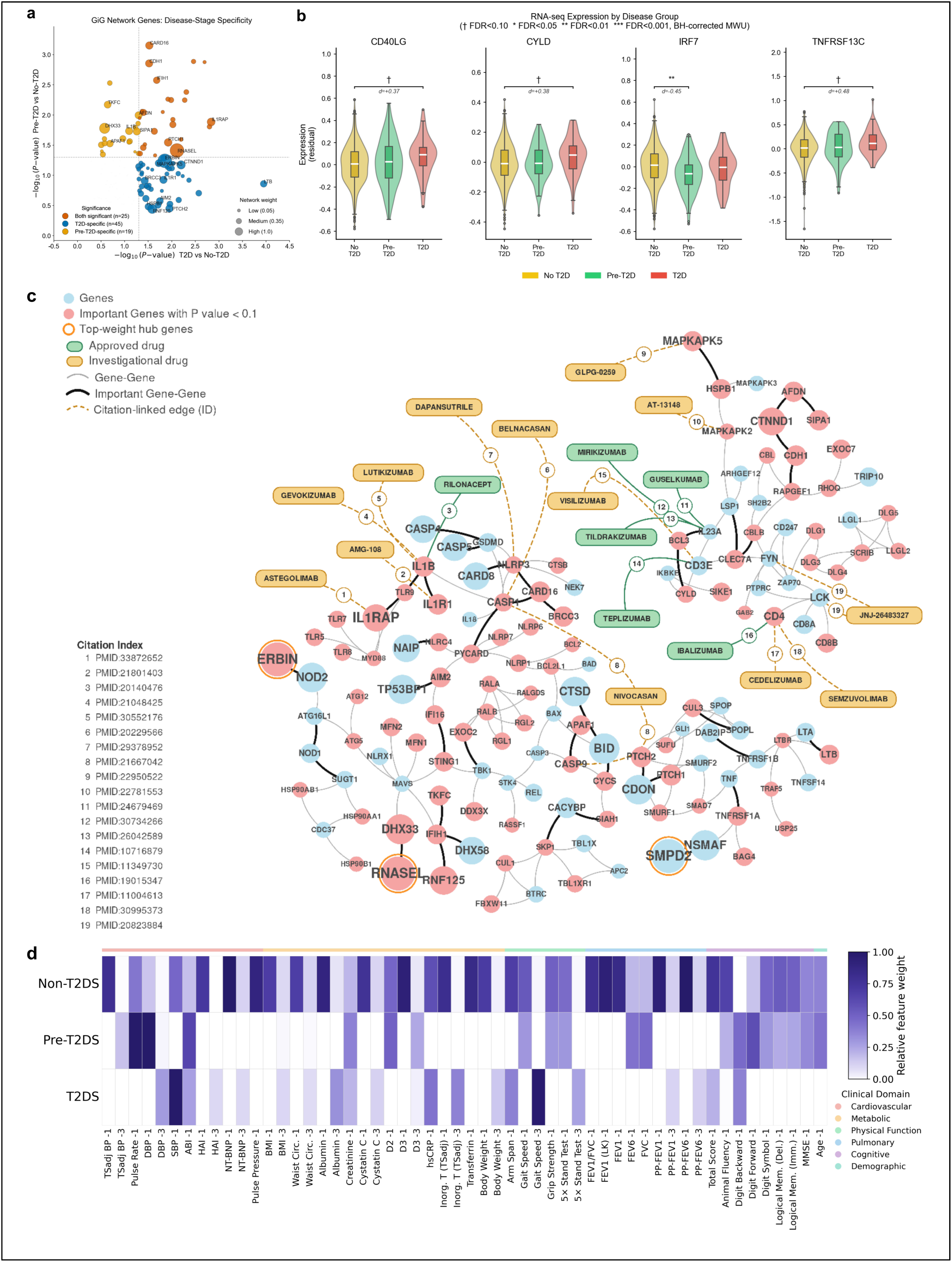
**a.** Disease-stage specificity of GiG network genes across T2D progression, stratified by expression significance. **b.** Residual RNA-seq expression distributions of four representative high-weight genes across no T2D, pre-T2D, and T2D groups. **c.** Drug-augmented GiG core network of T2D-focused overlay. Validated target-drug evidence from the final drug-evidence ledger is projected onto the T2D-focused GiG core network derived from Figure 2a. The original gene-gene topology and node-significance encoding are preserved, while drug nodes and target-drug links are added as an overlay. Citation-linked edges are shown as dashed lines, and edge ID labels map to PubMed citations in the in-figure Citation Index. The final overlay contains 21 target-drug edges across 13 core-network targets and 19 drug nodes, with approved and investigational drugs displayed using distinct node styles. **d.** Top phenotypic biomarkers across t2ds, pre_t2ds, and no_t2ds subgroups, where the suffixes −1, −2, and −3 denote the bottom 5%, middle 90%, and top 5% percentile ranges of each feature group, respectively.

### 2.4 Drugs targeting on the key signaling targets

To evaluate whether GiG-prioritized genes connect to actionable pharmacologic space, we apply the drug-evidence augmentation workflow to all 142 prioritized targets and construct a validated, citation-auditable evidence ledger (Figure 3c). The final merged ledger contains 159 rows across 142 targets, including 42 validated non-placeholder target-drug rows and 117 standardized placeholder rows indicating that no direct validated evidence is retained under current retrieval and validation rules. At the target level, 25 of 142 genes show at least one validated non-placeholder linkage. A compact summary of retrieval coverage, validation outcomes, and development-status distribution is provided in Table 2a.

**Table 2.**
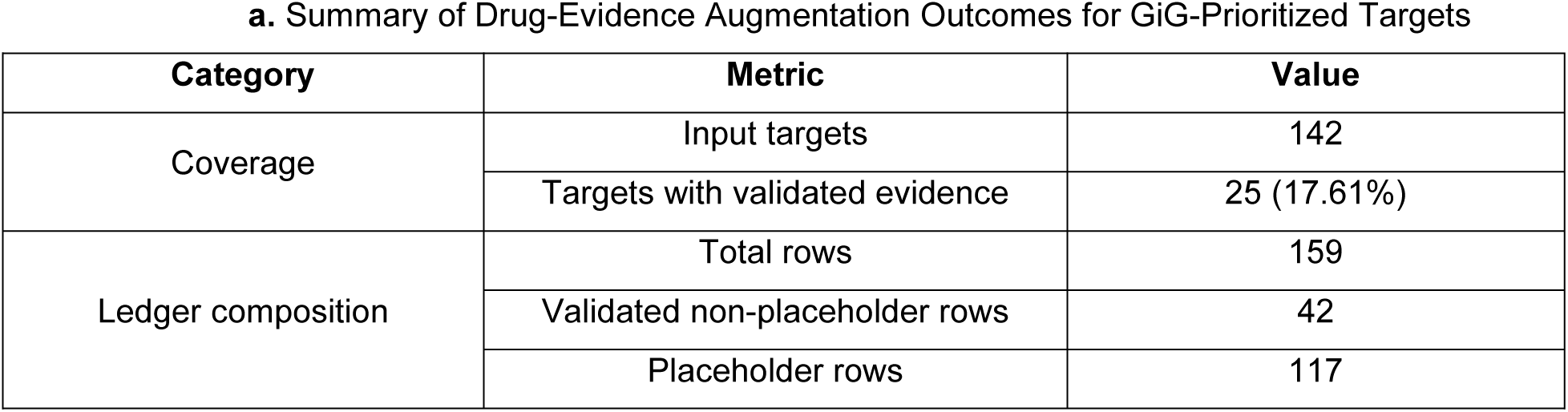
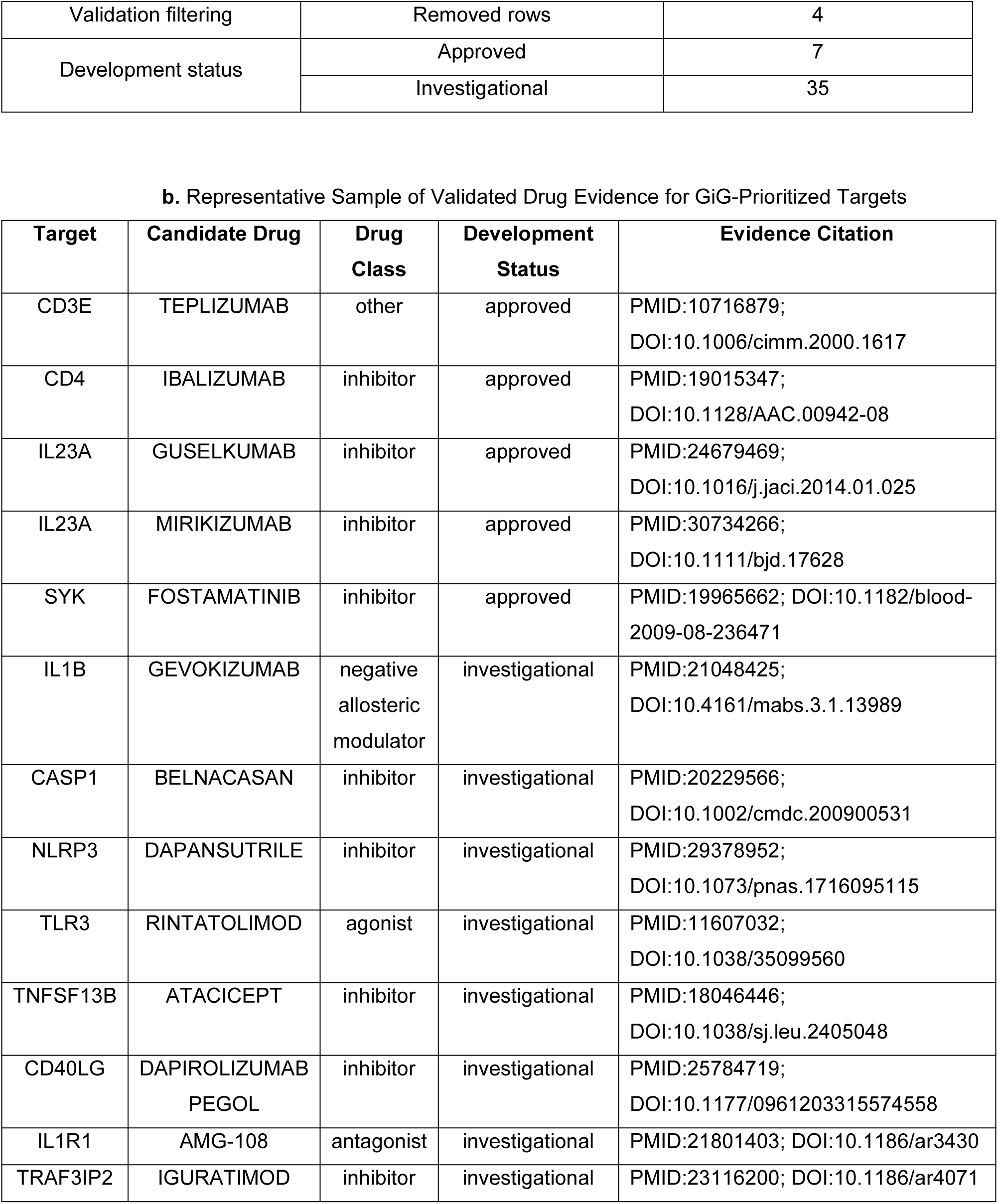
Drug-evidence augmentation results for GiG-prioritized targets. (a) summarizes coverage, validationfiltering, development-status distribution, and T2D-focused network mapping of the post-GiG drug-evidence workflow. (b) provides representative validated target-drug-citation examples from the final non-placeholder evidence ledger.

Among validated non-placeholder rows, development status includes 7 approved and 35 investigational rows, with no retained preclinical-only rows after confidence filtering. Validation accounting further reflects the conservative behavior of the pipeline: 42 candidate rows are retained as validated, 4 rows are removed during citation-consistency checks, and 117 targets are represented by fallback placeholder rows. This design emphasizes traceable support and reduces the risk of overstating druggability for genes without direct mechanism-linked evidence.

Approved-linked evidence is concentrated in immuno-inflammatory targets, including CD3E, CD4, IL1B, IL23A, and SYK. The broader investigational landscape additionally includes CASP1, CASP8/9, CD40/CD40LG, IL1R1/IL1RAP, NLRP3, TLR3, TNFRSF13C, TNFSF13B, and TRAF3IP2. This profile is consistent with the inflammatory and innate-immune emphasis of the GiG-derived network, indicating that high-weight network biology is partially connected to ongoing therapeutic programs.

We then project validated target-drug evidence onto the T2D-focused core network (Figure 2f) to generate a drug-augmented overlay (Figure 3c). In this overlay, 21 target-drug edges are mapped across 13 core-network targets and 19 drug nodes, enabling direct visual inspection of where translationally supported targets reside in the core signaling architecture. Together with Table 2, this figure provides a concise translational readout that links mechanistic prioritization to auditable intervention hypotheses.

Interpretation remains conservative. Target-level evidence mapping does not establish efficacy in the T2D/pre-T2D indication context, and development status reflects global drug-development context rather than T2D-specific approval. In addition, absence of validated direct evidence in this ledger should not be interpreted as absence of therapeutic potential, because it may reflect curation density, citation granularity, or strict confidence filters. Overall, this section extends GiG from mechanistic prioritization to a translationally annotated network output that is auditable and suitable for downstream therapeutic triage.

### 2.5 Key phenotypic biomarkers

By analyzing the group features with suffixes, we identified distinct phenotypic biomarker profiles for the three patient groups.The group features with suffixes indicate the range of subfeatures in each domain. For example, the group feature ‘_Pulsez_v1-1’ with suffix 1 means that this subfeature falls into the lowest five percentile of the whole feature. Suffix 2 implies that this subfeature falls into the range of the 5 to 95 percentiles of the entire feature group. Thus, the suffix 3 means that his subfeatures are over 95 percentiles of the whole feature group. Figure 3d displays the top subfeatures contributing most to the t2ds, pre_t2ds, and no_t2ds patient groups. Tables 3-5 detail the top 20 subfeatures for each of these categories.

**Table 3.** Top 20 features for t2ds patients (barplot)

| Rank | Group Name | Subgroup Name | Weight |
| --- | --- | --- | --- |
| 1 | SBP | SBP-1 | 0.0667 |
| 2 | Gait Speed | Gait Speed-3 | 0.0645 |
| 3 | hsCRP | hsCRP-1 | 0.0286 |
| 4 | Arm Span | Arm Span-1 | 0.0250 |
| 5 | Digit Backward | Digit Backward-1 | 0.0250 |
| 6 | Inorg. T (TSadj) | Inorg. T (TSadj)-3 | 0.0247 |
| 7 | Albumin | Albumin-3 | 0.0212 |
| 8 | DBP | DBP-3 | 0.0191 |
| 9 | ABI | ABI-1 | 0.0180 |
| 10 | 5x Stand Test | 5x Stand Test-3 | 0.0173 |
| 11 | Total Score (SPPB) | Total Score (SPPB)-1 | 0.0143 |
| 12 | Creatinine | Creatinine-1 | 0.0133 |
| 13 | Body Weight | Body Weight-3 | 0.0100 |
| 14 | PP-FEV6 | PP-FEV6-3 | 0.0084 |
| 15 | Cystatin C | Cystatin C-3 | 0.0078 |
| 16 | PP-FEV1 | PP-FEV1-3 | 0.0072 |
| 17 | BMI | BMI-3 | 0.0066 |
| 18 | HAI | HAI-3 | 0.0062 |
| 19 | NT-BNP | NT-BNP-3 | 0.0061 |
| 20 | Waist Circ. | Waist Circ.-3 | 0.0055 |

**Table 4.** Top 20 features for pre_t2ds patients.

| Rank | Group Name | Subgroup Name | Weight |
| --- | --- | --- | --- |
| 1 | Pulse Rate | Pulse Rate-1 | 0.1333 |
| 2 | DBP | DBP-1 | 0.1333 |
| 3 | ABI | ABI-1 | 0.0857 |
| 4 | Digit Forward | Digit Forward-1 | 0.0800 |
| 5 | D2 | D2-1 | 0.0667 |
| 6 | Digit Backward | Digit Backward-1 | 0.0615 |
| 7 | FVC | FVC-1 | 0.0615 |
| 8 | Age | Age-1 | 0.0571 |
| 9 | FEV6 | FEV6-1 | 0.0571 |
| 10 | Creatinine | Creatinine-1 | 0.0500 |
| 11 | MMSE | MMSE-1 | 0.0462 |
| 12 | Gait Speed | Gait Speed-1 | 0.0401 |
| 13 | Digit Symbol | Digit Symbol-1 | 0.0400 |
| 14 | Grip Strength | Grip Strength-1 | 0.0400 |
| 15 | 5x Stand Test | 5x Stand Test-1 | 0.0399 |
| 16 | Logical Mem. (Imm.) | Logical Mem. (Imm.)-1 | 0.0333 |
| 17 | D3 | D3-3 | 0.0324 |
| 18 | Logical Mem. (Del.) | Logical Mem. (Del.)-1 | 0.0308 |
| 19 | Animal Fluency | Animal Fluency-1 | 0.0286 |
| 20 | TSadj BP | TSadj BP-3 | 0.0220 |

**Table 5.**
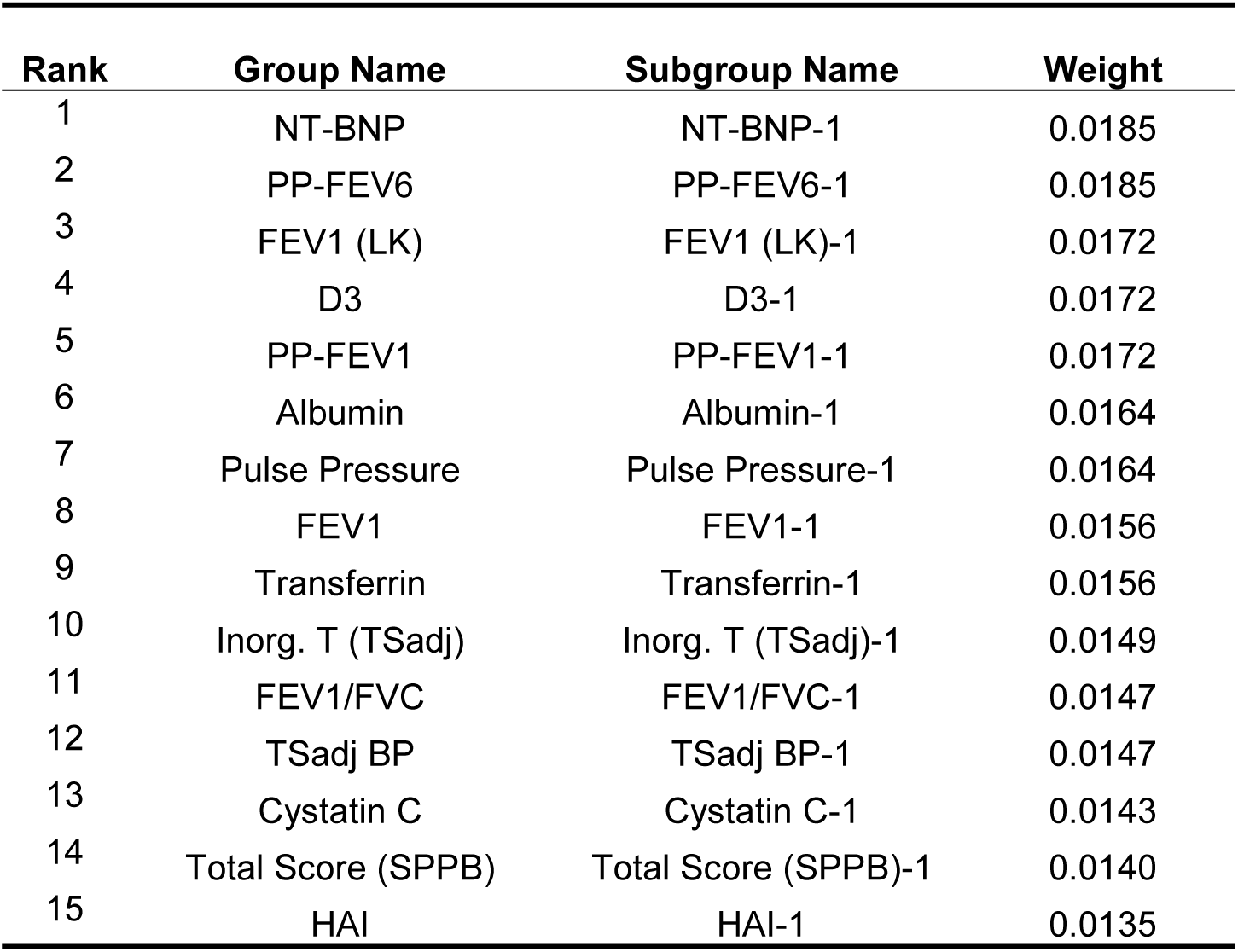
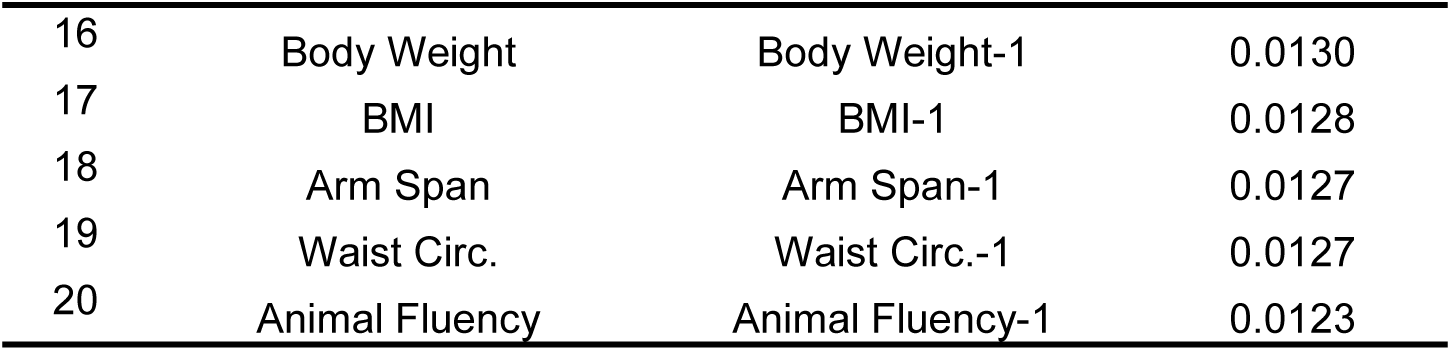
Top 20 features for no_t2ds patients.

### 2.6 Pre-T2D and T2D pathway analysis

These categorizations are derived from the high-level classifications provided in sources such as KEGG PATHWAY database^56^ and Gene Ontology (GO) Biological Process (BP)^57,58^.which broadly group pathways into categories like metabolism, genetic information processing, environmental information processing, cellular processes, organismal systems, human diseases, and drug development. The above categorizations are structured with a more specific focus on the functional or disease-related aspects of the pathways (Table 6-7).

**Table 6.**
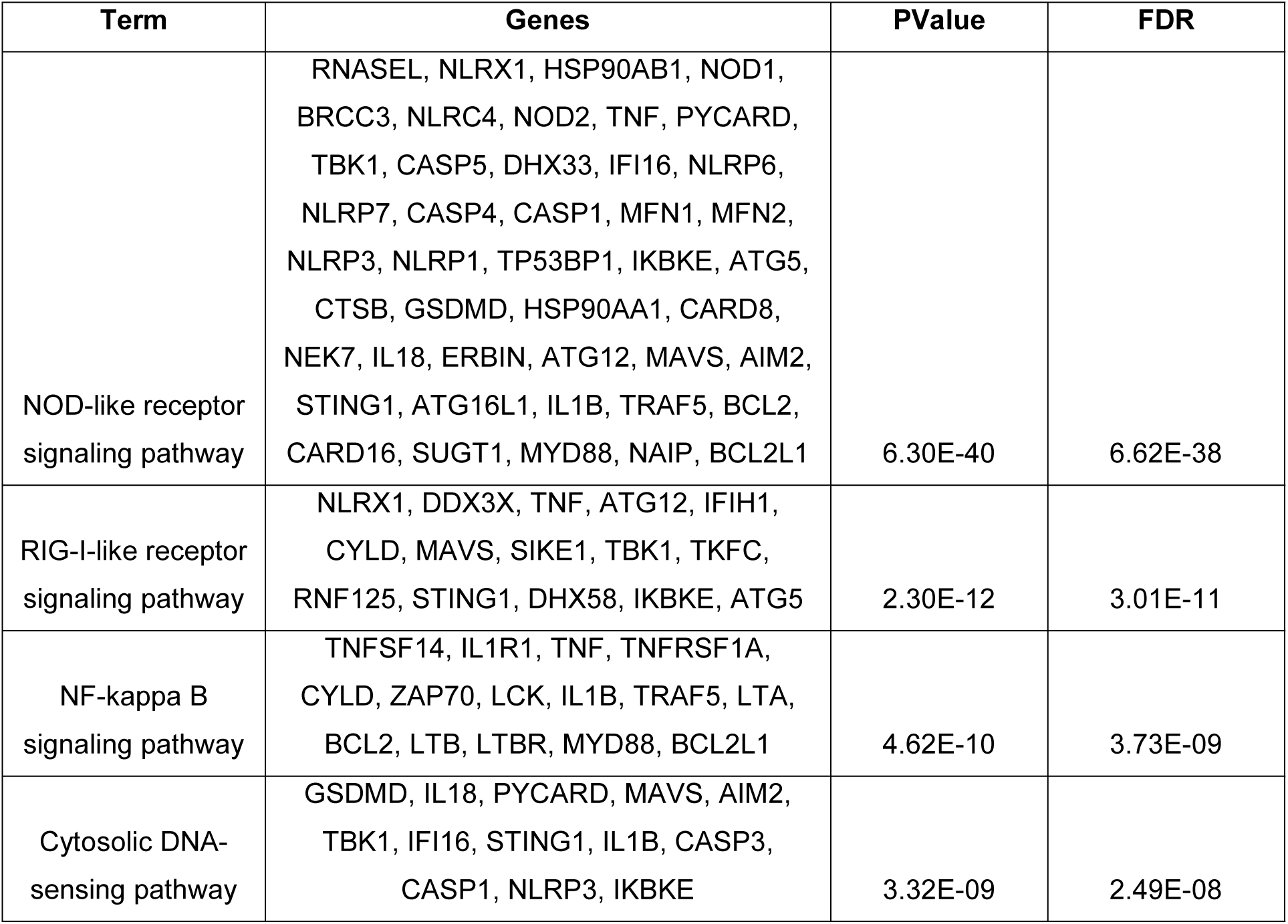
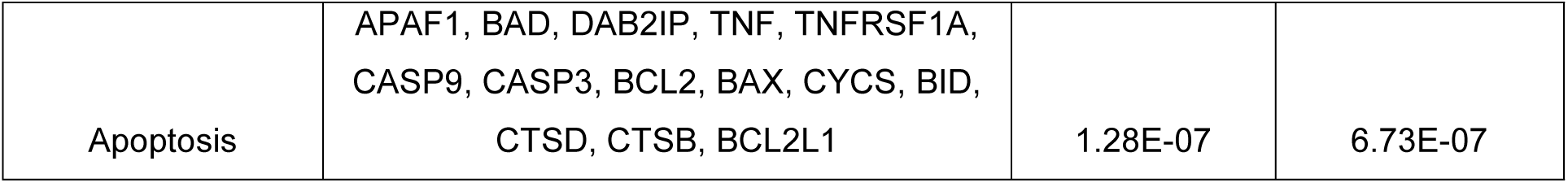
KEGG Pathway enrichment analysis.

**Table 7.**
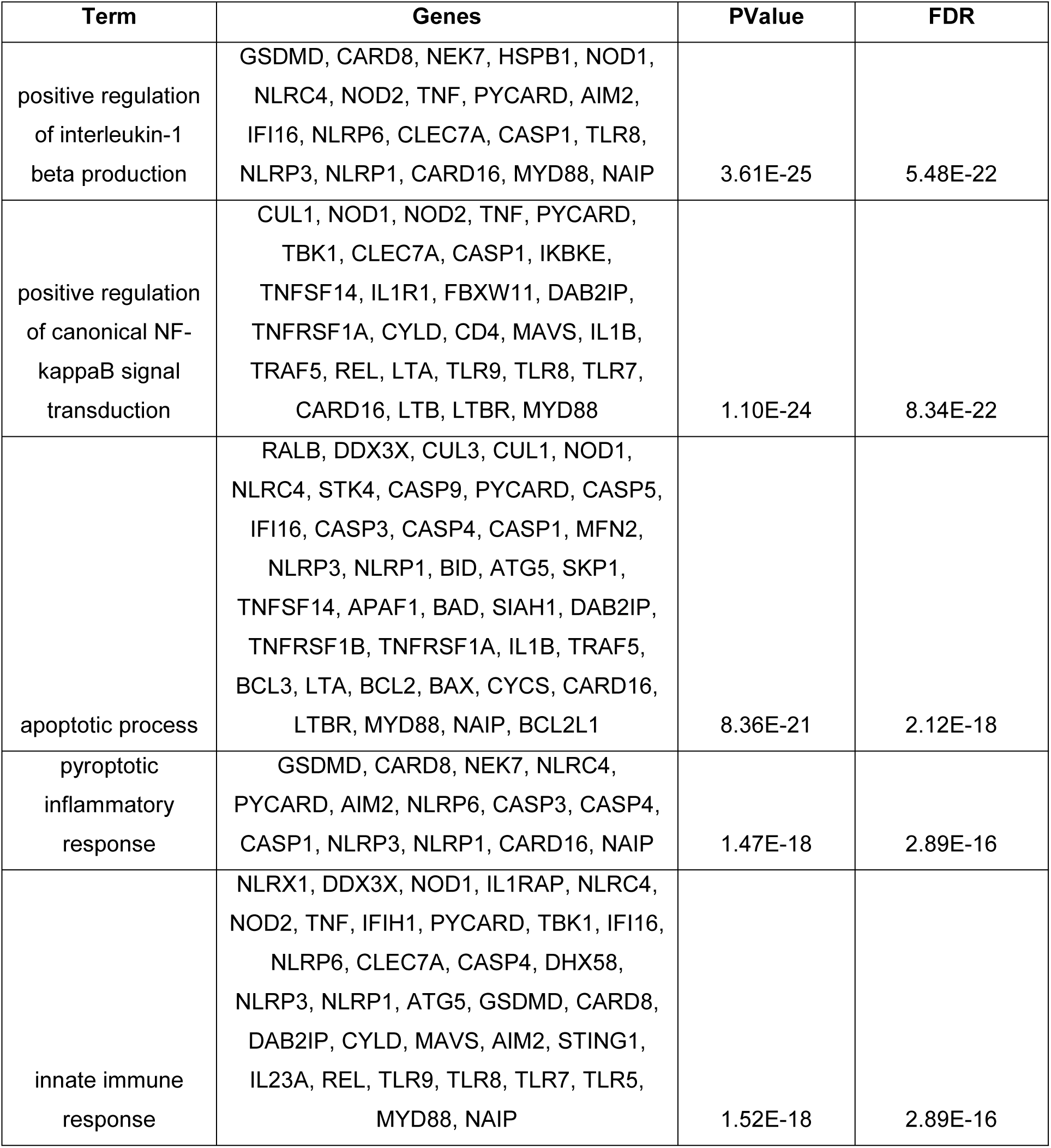
GO Pathway enrichment analysis.

As summarized in **Figure 4a**, pathway and ontology enrichment analyses across multiple annotation databases consistently converge on innate immune signaling and inflammasome-mediated cell death as the dominant biological themes. Among the KEGG pathways, the RIG-I-like receptor signaling pathway achieves the second-highest fold enrichment, followed by the NOD-like receptor signaling pathway and the NF-B signaling pathway, which rank third and fourth, respectively. This indicates that the prioritized genes are disproportionately concentrated in innate immune sensing and cytokine-driven inflammatory cascades. At the GO Biological Process level, immune system process and innate immune response rank as the most significantly enriched terms, with pattern recognition receptor signaling pathway exhibiting the largest fold enrichment among all GO BP terms. Pyroptotic inflammatory response and positive regulation of interleukin-1 beta production are also prominently enriched, directly implicating the inflammasome-caspase axis^59^ in the biological signal captured by the model. GO Molecular Function analysis reveals strong enrichment for CARD domain binding and cysteine-type endopeptidase activator activity, reflecting the functional specificity of the gene set toward inflammasome assembly interfaces and caspase catalytic regulation. At the cellular compartment level, canonical inflammasome complex and NLRP3 inflammasome complex display the most extreme fold enrichment values, with NLRP1 inflammasome complex and ripoptosome also significantly over-represented, collectively pointing to the intracellular multi-protein death-signaling machinery as the spatial substrate of the enrichment signal.

**Figure 4.**
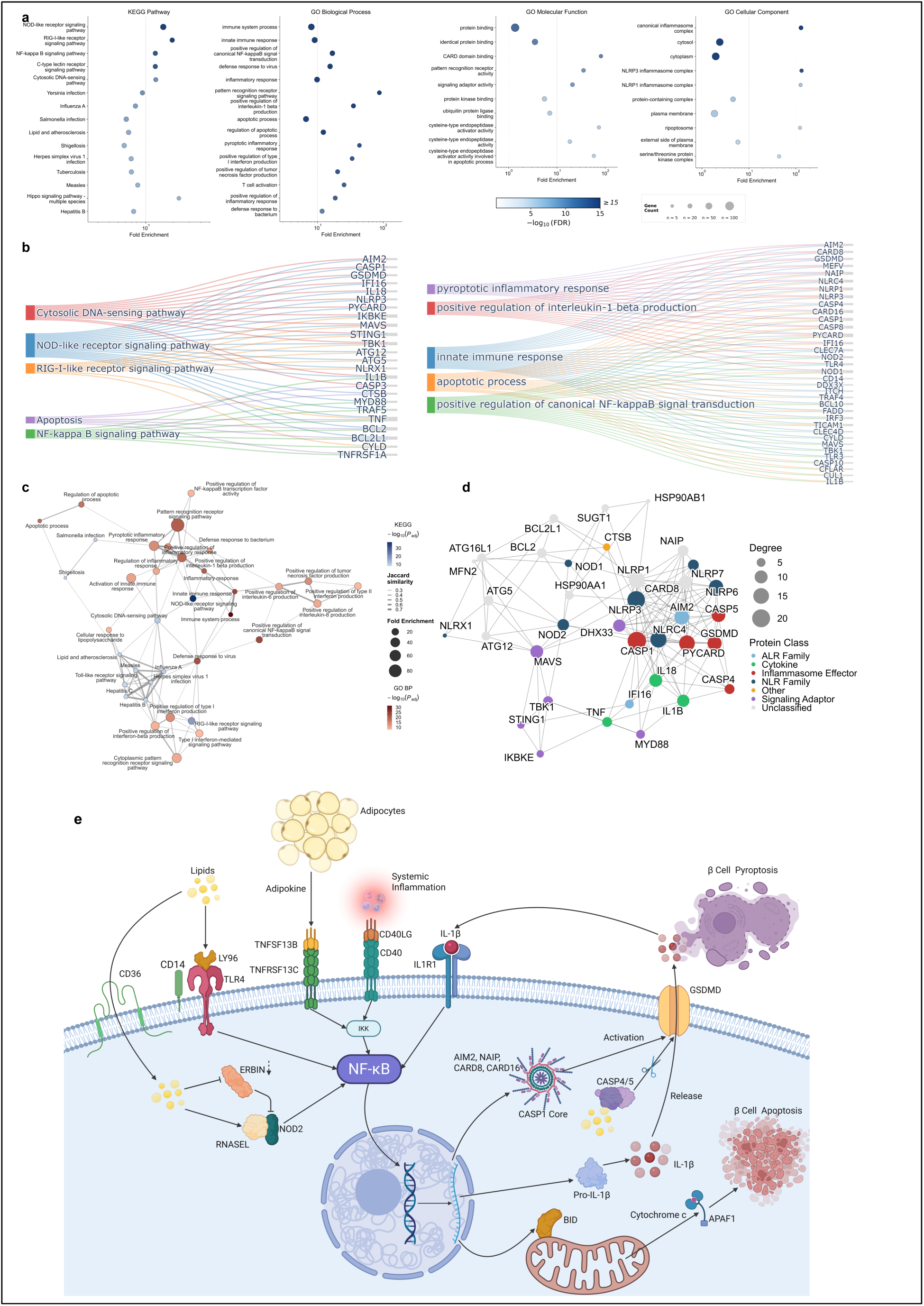
**a.** Functional enrichment analysis of prioritized genes across KEGG pathways, GO biological process, GO molecular function, and GO cellular component, highlighting prominent over-representation in innate immune signaling, inflammasome assembly, and caspase regulatory pathways. **b.** P-values and FDRs of top five representative GO terms and KEGG pathways. **c.** Enrichment map of pathway co-enrichment network, with nodes representing individual KEGG pathways and GO BP. **d.** Protein-protein interaction network of NOD-related inflammasome components, with node size scaled by interaction degree and color indicating protein class. **e.** Molecular mechanisms of β-cell death via NOD-like receptor–NF-κB–inflammasome signaling in the diabetic microenvironment.

To delineate the gene-level composition of the most enriched pathways, we constructed a Sankey diagram mapping individual genes to their respective pathway assignments (Figure 4b). To visualize the topological organization of these enriched terms and assess their interdependence, we constructed an enrichment map in which nodes represent individual pathways and edge weights reflect pairwise Jaccard similarity of their gene membership (Figure 4c). The resulting network resolves into two principal connectivity regions. The upper region is anchored by pattern recognition receptor signaling pathway, the highest-degree hub bridging inflammatory regulation terms (e.g., positive regulation of NF-κB, pyroptotic response, IL-1β production, and inflammatory response). The lower region is organized around antiviral innate immunity, where RIG-I-like receptor signaling pathway, type I interferon-mediated signaling pathway^60^, and cytoplasmic pattern recognition receptor signaling pathway^61^ form a tightly interconnected subcluster connected to viral infection pathways including Influenza A, Hepatitis B, Hepatitis C, and Herpes simplex virus 1 infection. The two regions are bridged through shared gene membership in innate immune response and cytosolic DNA-sensing pathway^62^, suggesting that the transcriptional program captured by the model integrates both sterile inflammatory and antiviral defense components into a unified regulatory structure. Apoptotic process and regulation of apoptotic process form a peripheral module linked to the inflammatory core, consistent with the role of caspase-mediated apoptosis^63^ as a functionally adjacent yet mechanistically distinct outcome of the same upstream danger-sensing machinery.

To further dissect the molecular architecture underlying the enriched inflammatory pathways, we constructed a protein-protein interaction (PPI) network centered on NOD-related inflammasome components (Figure 4d). Nodes are scaled by interaction degree and colored by protein class, encompassing NLR family members (NOD1, NOD2, NLRP1, NLRP3, NLRP6, NLRP7, NLRC4), inflammasome effectors (CASP1, CASP4, CASP5, GSDMD, PYCARD), ALR family sensors (AIM2, IFI16), cytokines (IL1B, IL18, TNF), and signaling adaptors (MAVS, MYD88, STING1, TBK1, IKBKE). CASP1 and PYCARD emerge as the highest-degree hubs, consistent with their central roles in inflammasome assembly and effector activation^59,64^, while GSDMD bridges this core complex to pyroptotic membrane execution^65,66^. The network further reveals functional crosstalk between canonical NOD/NLRP3 signaling and the cGAS-STING innate immune axis^61^ through MAVS, TBK1, and STING1, as well as between inflammasome activation and autophagy regulators including ATG5, ATG12, and ATG16L1^67^. These topological features indicate that the pathway enrichment signals identified above reflect a convergent, rather than modular, inflammatory architecture operating in the diabetic context.

### 2.7 Pathophysiological implications of the identified pathways

#### Cell Death Signaling in Diabetic β-Cell Dysfunction

The diabetic microenvironment introduces metabolic toxicity signals^68^ that are initially detected by intracellular NOD-like receptor pathways^69^ where the heavily weighted genes ERBIN^70^, RNASEL^42^, and NOD2^70^ function as frontline danger sensors (Figure 4e, left panel). ERBIN acts as a critical regulator of NOD2^70^, and the activation of this receptor directly induces cell-autonomous innate immune responses^71^ that initiate inflammatory signaling and contribute to insulin resistance^70^. These sensory signals rapidly propagate to hyperactivate the NF-κB signaling hub through mediators such as the adipokine TNFSF13B (Figure 4e, center panel)^72^, which is abundantly secreted by adipose tissue under obese conditions to engage its receptor TNFRSF13C and stimulate the IKK complex^73^. The resulting dysregulation of NF-κB, exacerbated by ERBIN deficiency, drives a systemic chronic inflammatory cascade characterized by the substantial production of pro-inflammatory cytokines including IL-1 β ^42,74^. This severe inflammatory stress ultimately recruits downstream executioner proteins, specifically the model-identified components CASP4^65^ and BID (Figure 4e, right panel)^75^, to drive irreversible programmed cell death. This terminal phase comprises apoptosis as well as parallel inflammatory modalities including pyroptosis and necroptosis, which directly cause the rupture and death of pancreatic β-cells. The prominence of these mechanisms is robustly supported by the functional enrichment of specific biological processes, particularly pyroptotic cell death, pyroptosome complex assembly, and the positive regulation of the pyroptotic inflammatory response, alongside the necroptotic signaling pathway and the general regulation of the necroptotic process. The subsequent maturation and release of IL-1β engage IL1R1 to establish a destructive auto-amplification loop^76,77^, demonstrating that this integrated network of programmed cell death pathways operates as a central pathogenic driver rather than a peripheral inflammatory byproduct in the disease progression^78^.

#### Lipid Metabolism and Cardiovascular Complications

Beyond direct cellular toxicity, the systemic metabolic dysregulation in T2D profoundly alters lipid homeostasis, establishing a critical mechanistic link to cardiovascular complications^51^. This pathogenic axis is strongly supported by the specific enrichment of the lipid and atherosclerosis pathway in our analysis. The functional integration of these enrichment terms reveals a robust intersection between aberrant lipid metabolism and the innate immune networks. Specifically, the accumulation of metabolic byproducts, notably oxidized low-density lipoproteins, acts as a potent stimulus that directly engages pattern recognition receptors^79^. This cellular response to oxidized lipids subsequently triggers the assembly and positive regulation of the NLRP3 inflammasome while simultaneously driving the canonical NF-κ B signaling cascade^80^. Within the vascular microenvironment, this continuous inflammatory signaling exacerbates macrophage activation and foam cell formation, thereby accelerating atherosclerotic progression^81^. Consequently, these findings indicate that the lipid-driven inflammatory feedback loop is not merely a secondary manifestation of diabetes but a fundamental driver that mechanistically bridges metabolic dysfunction with severe cardiovascular comorbidities.

#### Immunological Disorders

The pathways related to Inflammatory Bowel Disease (IBD), Rheumatoid Arthritis (RA), and the Intestinal Immune Network for IgA Production, have interconnected aspects that might have implications in the context of Type 2 Diabetes (T2D)^82^. Recent findings suggest a notable link between the metabolites produced by gut microbes and the subsequent inflammation in the intestinal and systemic regions, with the onset and advancement of diabetes^83^. It’s estimated that nearly 90% of type 2 diabetes cases are connected to imbalances in the gut microbiota, known as dysbiosis. This kind of microbial imbalance is also observed in cases of Inflammatory Bowel Disease (IBD)^84,85^. Aside from that, there’s a pathogenic connection between RA and T2D, forming a vicious circle driven by glucose derangement and inflammatory mediators. This suggests that the immunological disturbances in RA could have a cascading effect, leading to or worsening T2D ^86^. RA is associated with anomalies in glucose metabolism, primarily insulin resistance, which could progress to T2D. This association suggests a metabolic and immunological crosstalk between the two conditions^87^. And a retrospective cohort study found that there could be shared pathogenesis stemming from the shared genetics of immune-related diseases between T2D and RA, indicating a possible genetic and immunological basis for the association between these conditions^88^. What’s more, some research has hinted at novel regulatory pathways within the intestinal immune network for IgA production, which could be relevant in the context of T2D. For instance, an investigation in black rockfish revealed a potential regulatory pathway of the immunity along the intestine-spleen axis, which might be of interest in understanding the immunometabolic interactions in T2D^89^.

#### Bacterial infections

Tuberculosis and Salmonella infection pathways are highly related to type 2 diabetes. Salmonella can cause extra-intestinal focal infections, apart from the common gastrointestinal problems. In some documented cases, individuals with T2D, especially those with poor glycemic control, have been found to be susceptible to severe Salmonella infections. For instance, a case study reported a 30-year-old man with poorly controlled T2D who developed a severe Salmonella skin and soft tissue infection leading to a chest wall abscess. This infection required surgical intervention and the patient was treated with antibiotics alongside insulin therapy to manage his blood sugar levels^90^. In another case, individuals with Type 2 Diabetes are at an increased risk for infections, including the development of abscesses, due to impaired immune responses and altered tissue conditions. High blood sugar levels can impair the function of immune cells, slow down blood circulation, and affect the body’s ability to heal, which makes it easier for infections to take hold and develop into abscesses. An abscess, which is a collection of pus that has formed within the tissue, can occur at different sites in the body, and its management can be complicated in individuals with diabetes, requiring meticulous glycemic control alongside infection management^91^. Furthermore, diabetes impairs the immune system, making it more challenging for the body to combat TB bacteria^92^. And the evidence shows that diabetes increases the likelihood of developing active TB. Individuals with diabetes have about a two to three-fold higher risk of developing TB compared to those without diabetes^93^.

## 3 Discussion

In this study, for the first time, we introduced Graph in Graph (GiG), which is a novel dual-graph AI model and is capable of systematically integrating and interpreting complex medical record phenotypes and high-dimensional omic datasets at the individual patient level. While the availability of clinical records and biological omics has created new avenues for precision healthcare, effective multi-modal integration has remained a significant computational bottleneck. GiG mitigates these challenges by leveraging a dual-graph structure: a person-phenotype graph to capture clinical features alongside patient-specific omics signaling graphs. This design enables seamless representation across the whole-person biological spectrum, bridging the gap between molecular mechanisms and macro-level clinical manifestations.

Applying GiG to the Long Life Family Study (LLFS) cohort demonstrated both strong predictive performance and high model interpretability in evaluating Type 2 Diabetes (T2D), pre-T2D, and control profiles. The model uncovered key clinical phenotypes and multi-omic biomarkers associated with disease progression. Beyond candidate biomarker identification, GiG successfully pinpointed critical signaling pathways driving metabolic dysfunction and prioritized potential therapeutic drugs, offering valuable mechanistically backed hypotheses for translational research. Overall, the GiG framework provides a scalable, generalizable AI framework for precision medicine. By effectively synthesizing heterogeneous biological and clinical data to drive both diagnosis and biomarker discovery, GiG can be readily adapted to investigate diverse disease contexts with both medical records and omics datasets.

## 4. Methodology

### 4.1 Study Population Design

This study is based on multi-omics and clinical phenotype data from the Long Life Family Study (LLFS) cohort. Transcriptomic profiles were available for 1,810 subjects, DNA methylation data (Illumina Methylation450K array, summarized across five gene-regulatory regions—core promoter, proximal promoter, distal promoter, upstream, and downstream) were available for 936 subjects, and clinical phenotype data were available for 4,810 subjects. Subjects were retained in the final analytic sample only if they had complete data across all three modalities— transcriptomics, DNA methylation, and clinical phenotype, including type 2 diabetes (T2D) status—yielding a final cohort of 813 subjects.

Subjects were stratified into three mutually exclusive groups based on established LLFS T2D status classifications: T2D (n=68), pre-T2D (n=104), and No-T2D (n=641). This three-group T2D status served both as the between-group comparison variable and as the prediction target (classification label) for the graph neural network model.

In addition to T2D status, 34 continuous clinical phenotypes were incorporated as phenotype-channel features and used for group-wise statistical comparisons, spanning demographic (age), anthropometric (BMI, waist circumference, weight), cardiovascular (systolic/diastolic blood pressure z-scores, pulse pressure, pulse rate), metabolic (glucose, HbA1c, and lipid profile), pulmonary function (FEV1, FEV6, FEV1/FVC, FVC), cognitive function (MMSE total score, digit span, digit symbol substitution, logical memory, animal naming), and physical performance (gait speed, chair-stand time, grip strength) domains. All continuous phenotypes were z-score standardized, with log- or inverse-normal transformation applied as needed prior to normalization.

### 4.2 Omics Data

Three data types were used to construct the model. Transcriptomic data consisted of covariate-adjusted expression residuals for 16,419 genes (Ensembl gene IDs) across 1,810 subjects^94^. DNA methylation data, derived from the Illumina Methylation450K array, were organized as gene-level β-values (0–1) across the five regulatory regions described above, each covering 936 subjects. Clinical phenotype and T2D status labels were obtained from the LLFS phenotype database. The dataset was accessible at: https://prod.eliteportal.synapse.org/Explore/Projects/DetailsPage?shortName=LLFS

To prepare these data for machine learning, transcriptomic and methylation probes were first mapped to gene-level identifiers via Ensembl annotation, and the resulting gene sets were intersected across the transcriptomic, methylation, and KEGG pathway network gene lists, yielding a final panel of 1,390 genes (reduced from 16,419). Subject identifiers were then intersected across the phenotype, T2D-label, transcriptomic, and methylation datasets to obtain the final analytic sample of 813 subjects (T2D=68, pre-T2D=104, No-T2D=641). Phenotype data were cleaned and restricted to the visit-1 timepoint, one-hot encoded, and discretized into categorical bins using the 10th/90th percentile thresholds to construct subject–phenotype bipartite subfeature nodes.

Gene expression and methylation feature matrices were each normalized and concatenated into a six-channel gene-node feature representation (one transcriptomic channel plus five methylation-region channels); normalized phenotype features were concatenated along the subject dimension to form the final node feature matrix. Gene–gene edges were derived from the KEGG pathway database (1,390 gene nodes; 9,199 edges), and subject–phenotype bipartite edges were derived from the discretized phenotype categories (813 subject nodes, 122 phenotype subfeature nodes; 32,600 edges). For exploratory group comparisons, each gene- and CpG-region-level feature was tested across the three pairwise T2D-status comparisons (T2D vs. No-T2D, T2D vs. pre-T2D, pre-T2D vs. No-T2D) using the Mann–Whitney U test, with multiple-testing correction by the Benjamini–Hochberg procedure.

### 4.3 Model of Graph in Graph (GiG)

#### Problem formulation

To construct this graph neural network framework, we considered embedding the graph neural network as a node in another graph, which will be used as the giant graph in this framework. In this study, we embedded the transcriptomic graphs as nodes in the subject-feature graph. Hence, for the patient-feature graph, *G* = (*V, E*), where 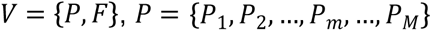, 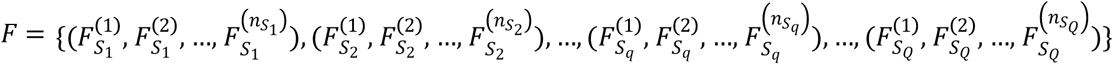 and 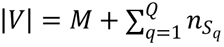. To be more specific, there are two types of nodes in the graph. Nodes set *Р* is the set containing patient nodes, and there are *M* patients in the graph *G*. And nodes set *F* is the set containing feature nodes, and there are *Q* groups of subfeatures. In certain subfeature groups *S_q_*, there are 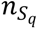 feature nodes. And we have patient phenotype feature matrix 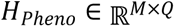 and subfeature matrix 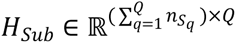. To implement the embedded graph in specific patient nodes *P_m_*, the embedded graph 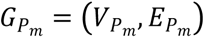 will be built, where 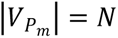 and all of 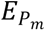 share the same graph structure with *E_g_*. In any of the embedded graph *P_m_*, the nodes set (i.e., the genes set) contains the transcriptomic data with node/gene feature matrix 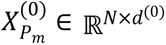, where *d*^(0)^ is the number of dimensions for the node initial feature. For the gene feature in the patient cohort, the gene feature matrix will be 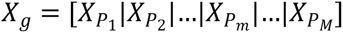, where 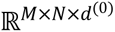.

Hence, the Graph in Graph model (GiG) will predict the type of patients for the cohort with the phenotype and genotype input 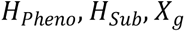,

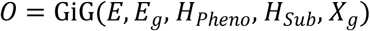

, where 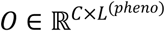 and *C* is the number of classes for the downstream task.

#### GiG Framework Architecture

The overall architecture of the GiG framework is illustrated in Figure 1a. To embed the graph as a feature vector for the node *P_m_* in the patient-feature graph, which will be the graphical model in the gene graph with 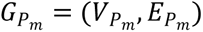, the attention-based model by graph transformer^95^ will be applied here for interpretation possibility through message passing over layers 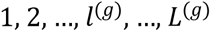:

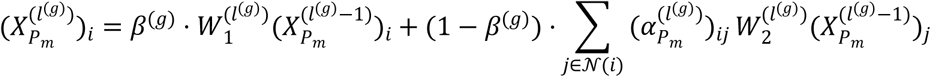

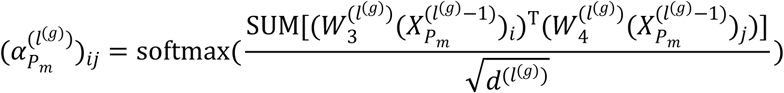

In the *l^(g)^* layer, the input node features is the set of *N* nodes/genes with 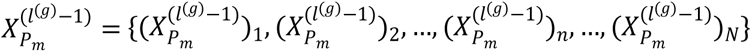, where 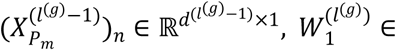 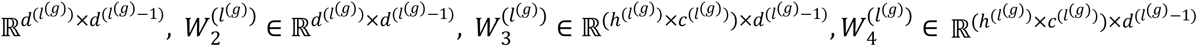 and there are *L*^(*g*)^ layers in total. 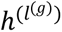 and 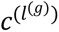 are the number of heads and channels in the *l^(g)^* layer. Here, the SUM function will aggregate all embedded features. Thus, the final attention coefficient between node *i* and node *j* will be 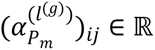 in the *l^(g)^* layer. To mitigate the dilution of attention weights over many features, we subsequently apply a threshold to prune low-weight edges and retain only the top-weighted nodes for interpretation.

With the embedded node features, 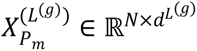, in the last layer, the pooling method based assignment matrix^96^ will be used for transforming the *N* nodes/genes into lower dimensions with *K* critical features by

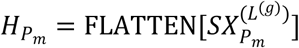

, where 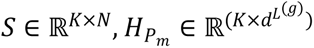.

After collecting the embedded graph from gene features of the individual patient, we can get the node/ patient features with 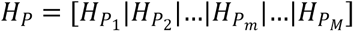, where 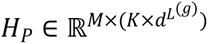 is the patient embedded genomic feature. From the initial patient-feature graph *G*, we have patient phenotype feature matrix 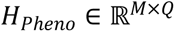 and subfeature matrix 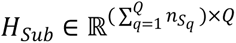 By concatenating the embedded genomic features and phenotype features for all patients, we have 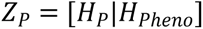, where 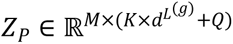. Leveraging the zero padding subfeature matrix,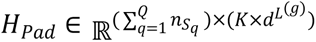, we will have subfeature concatenated matrix 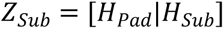, where 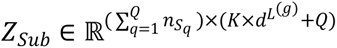. In this way, the initial feature of the patient-feature graph with concatenated feature matrix will be 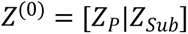, where 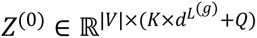. For convenience, we will use 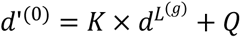.

In the end, the graph transformer will also be applied on the patient-feature graph message passing to embed the node features over layers 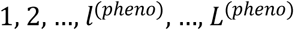. For certain layer *l^(pheno)^*, the message passing by the transformer will have the same procedure for the gene graph. **Finally,** we will have the embedded node features in the *L^(pheno)^* layer with 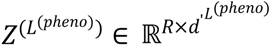. As we will make the node classification task, the linear transformation will be made by

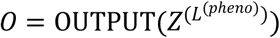

, where 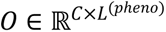 and *C* is the number of classes for the downstream task.

### 4.4 Experimental Settings and Classification Results

For the gene graph, we set the number of messages passing layer *L^(g)^* = 2. For the patient-feature graph, we set the number of messages passing layer *L^(pheno)^* =3. And for the rest of model parameters, you can check the **Table 8** for reference.

**Table 8.**
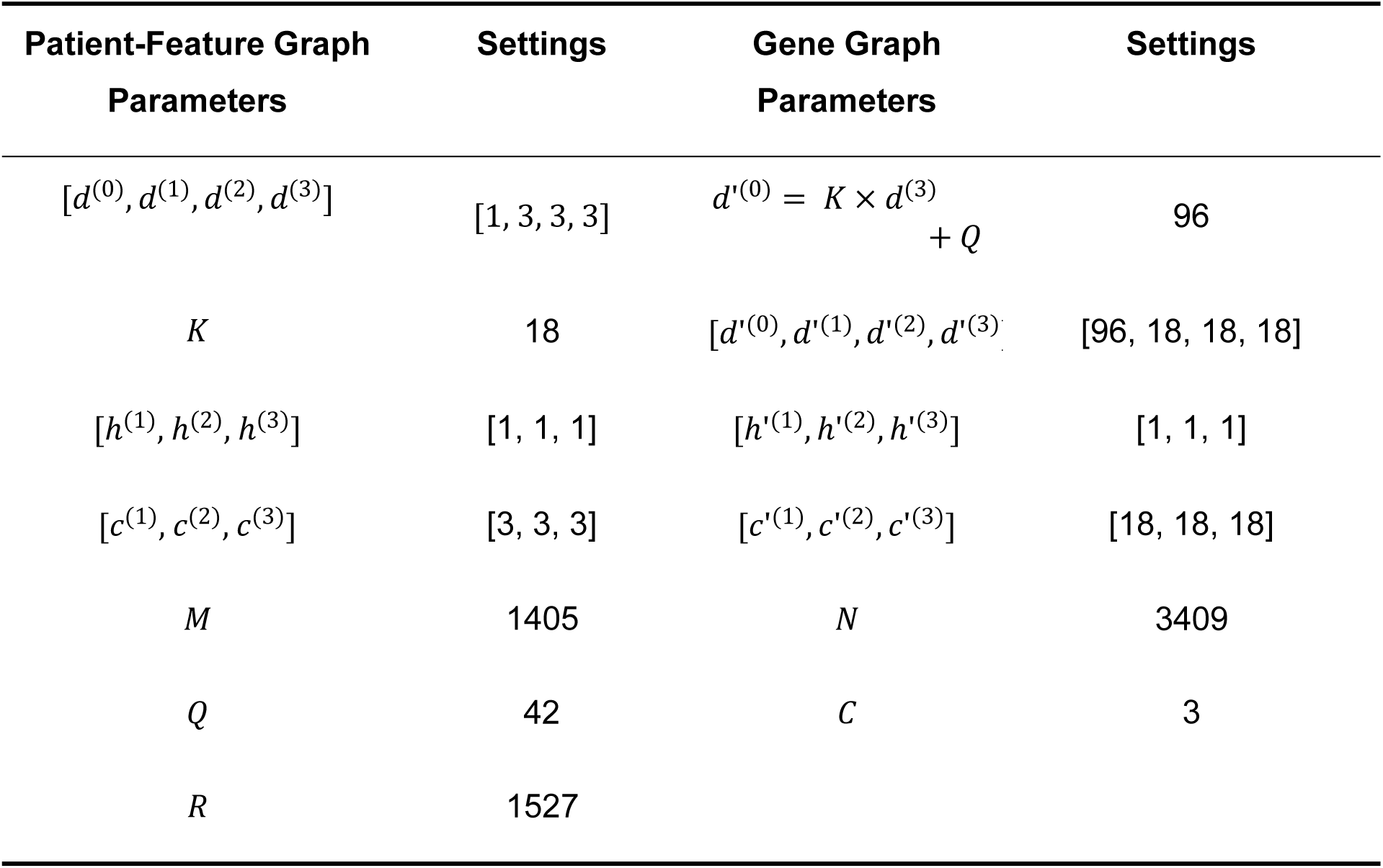
Model parameter and hyper-parameter settings.

Here, 5-fold cross-validation was used to run the experiments. Each used the same hyperparameters with the Adam optimizer, set with beta= [0.8, 0.999], eps=1e-7, weight_decay=1e-10. The multi-step learning rate schedule function in Pytorch was used by setting the stage with [100, 200, 300, 400, 500, 600, 700, 800, 900, 1000, 1250] and gamma=0.9 with learning rate starting from 0.01. To evaluate the contribution of each data modality and benchmark the dual-graph architecture against alternative GNN designs, we compared the full GiG model with two baseline architectures trained on the same combined input (GAT, GIN; Table 9). Across 5-fold cross-validation, the full GiG model achieved the highest mean test accuracy (80.57% ± 0.79%), outperforming the baseline architectures (GAT: 75.15% ± 2.84%; GIN: 77.25% ± 0.54%). These results indicate that jointly modeling phenotype and genotype within the GiG dual-graph architecture provides complementary predictive information beyond GNN architecture alone.

**Table 9.** Test prediction accuracy of GiG compared with baseline GNN architectures (GAT, GIN) for three-class T2D/pre-T2D/No-T2D classification on the LLFS dataset using 5-fold cross-validation.

| Data Split | GiG | GAT | GIN |
| --- | --- | --- | --- |
| Fold 1 | 81.48% | 76.54% | 76.54% |
| Fold 2 | 80.86% | 70.99% | 77.78% |
| Fold 3 | 80.25% | 73.46% | 77.16% |
| Fold 4 | 80.86% | 77.78% | 77.78% |
| Fold 5 | 79.39% | 76.97% | 76.97% |
| Average | **$80.57\% \pm 0.79\%$** | $75.15\% \pm 2.84\%$ | $77.25\% \pm 0.54\%$ |

### 4.5 Drug-Evidence Augmentation of Prioritized Genes

To improve the translational interpretability of the GiG-prioritized signaling network, we implement a downstream drug-evidence augmentation module that maps prioritized genes to target-level pharmacologic evidence. This module is a post-GiG layer: it does not modify model architecture, optimization, or classification performance, but adds a structured evidence layer to network-level mechanistic outputs.

The module takes as input the top 142 GiG-prioritized genes derived from the core T2D/pre-T2D network. In this input set, all target symbols are unique; therefore, no additional deduplication step is required in the manuscript workflow. Each target is propagated directly to evidence mining.

Evidence retrieval follows a rule-constrained, citation-auditable pipeline (Figure 1b). For each target, we query ChEMBL human-target records, mechanism records, and molecule annotations. Candidate target-drug links are retained only when mechanism-level records with traceable literature support are available. Linked PubMed records are then parsed to extract citation metadata (PMID/DOI). Development status is harmonized by a controlled vocabulary (approved, investigational, preclinical, unknown), with phase-based normalization when source annotations differ.

To minimize unsupported claims, evidence generation is separated from evidence validation. In the generation stage, candidate rows are assembled into a structured ledger with five fields: Target, Candidate Drug, Drug Class, Evidence Citation (must include PMID/DOI/Author-Year), and Development Status. In the validation stage, each non-placeholder row undergoes lightweight citation-consistency checks against PubMed metadata and mechanism context. Rows that fail confidence criteria are removed. Targets without remaining validated non-placeholder evidence are automatically reverted to a standardized placeholder row (“No validated drug“; “No direct evidence found“; status = unknown). This fallback policy prevents over-interpretation when direct evidence is insufficient.

The workflow is executed in batched mode to maintain deterministic parsing and auditable intermediate outputs. Reproducibility assets include batch-level draft ledgers, batch-level validated ledgers, row-level validation reports, and a merged final validated ledger with summary statistics. Raw API payloads (target/mechanism/molecule and PubMed XML) are cached for inspection. After ledger construction, validated target-drug rows are projected onto the GiG core network to create drug-augmented overlays while preserving original gene-gene topology and node-significance encoding. This projection supports direct interpretation of translationally supported targets within the inflammatory and innate-immune network architecture identified by GiG.

## Acknowledgment

We extend our gratitude to the Long Life Family Study consortium, including the participants, investigators, and administrative and clinical staff. This study was supported by NIA 4R33AG078799-02, NIA 1R21AG078799-01A1, NIA AG063893.

## Data Availability

The data supporting this study are available from the NIA Long Life Family Study via the Synapse platform. Access details and application procedures can be found at https://prod.eliteportal.synapse.org/Explore/Projects/DetailsPage?shortName=LLFS.

